# Perceptual rivalry with vibrotactile stimuli

**DOI:** 10.1101/2020.09.17.301358

**Authors:** Farzaneh Darki, James Rankin

**Author notes:** Correspondence concerning this article should be addressed to Farzaneh Darki, Department of Mathematics, College of Engineering, Mathematics & Physical Sciences, University of Exeter.

## Abstract

In perceptual rivalry, ambiguous sensory information leads to dynamic changes in the perceptual interpretation of fixed stimuli. This phenomenon occurs when participants receive sensory stimuli that support two or more distinct interpretations; this results in spontaneous alternations between possible perceptual interpretations. Perceptual rivalry has been widely studied across different sensory modalities including vision, audition and, to a limited extent, in the tactile domain. Common features of perceptual rivalry across various ambiguous visual and auditory paradigms characterise the randomness of switching times and their dependence on input strength manipulations (Levelt’s propositions). It is still unclear whether general characteristics of perceptual rivalry are preserved with tactile stimuli. This study aims to introduce a simple tactile stimulus capable of generating perceptual rivalry and explores whether general features of perceptual rivalry from other modalities extend to the tactile domain.

---

Multistable perception occurs when sensory information is ambiguous and consistent with two or more perceptual states. In this phenomenon, perception alternates intermittently between two (bistable) or more (multistable) interpretations of the fixed stimulus [Sterzer et al., 2009]. Examples of multistable perception are well-known in vision such as motion direction with plaids (ambiguous barber poles) [Hupé and Rubin, 2003, Wuerger et al., 1996], apparent motion [Meso et al., 2016, Ramachandran and Anstis, 1983], and binocular rivalry [Blake, 1989], and span across other sensory modalities including audition [Pressnitzer and Hupé, 2006] and olfaction [Zhou and Chen, 2009]. All of these multistable phenomena share common features, such as exclusivity of perceptual interpretations, randomness, inevitability of alternations, independence of perceptual phases [Lathrop, 1966, Fox and Herrmann, 1967], and Levelt’s propositions [Levelt, 1968, Leopold and Logothetis, 1999]. These similar characteristics give rise to the conclusion that perceptual ambiguity must have a common neural basis and is likely resolved at a higher level of cognition that is not specific to individual sensory modalities [Pressnitzer and Hupé, 2006, Logothetis et al., 1996a, Wolfe, 1996]. A more recent study suggests that perceptual switching arises from a distributed system of similar but independent processes [Denham et al., 2018].

Many of the early studies of tactile illusions were focused on investigating the existence of tactual equivalents of apparent motion. These kind of illusory experiments were generally extensions of known illusions from vision [Lederman and Klatzky, 2009]. In visual experiments two spatially separated lights flash on and off in turn and can produce the illusion of movement (also known as phi phenomenon or beta movement). A tactile variant of smooth apparent motion was first produced with stimulation of two vibrotactile bursts of 150 Hz presented sequentially on the skin [Boring, 1942, Sherrick and Rogers, 1966]. Observers typically report that the series of discrete tactile vibrations feel like vibratory stimulation moving across the skin [Burtt, 1917, Kirman, 1974, Lederman and Jones, 2011, Burtt, 1917]. Various stimulus parameters such as the inter-stimulus onset interval (ISOI) and stimulus duration (from 25 to 400 ms) have been studied to find the optimal ranges for the induction of smooth apparent movement [Sherrick and Rogers, 1966]. Apparent motion can also happen with a bilateral stimulus delivered to each arm, however, the movement is less robust than the movement reported when stimulation is delivered to a unilateral limb such as a single thigh or a single arm [Sherrick, 1968].

Tactile pulses separated in location and time can induce a sensation of movement. For more complicated patterns, where different directions of movement are consistent with pulses, perception can be bistable. The first example of an apparent motion stimulus that leads to perception of motion in different directions is the “apparent motion quartet” stimulus realized in both the visual and tactile domains [Carter et al., 2008]. In different studies, pairs of vibrotactile stimuli were attached to the tip of a participant’s index finger [Carter et al., 2008], at locations on both index fingers [Conrad et al., 2012], to the thumbs and index fingers [Haladjian et al., 2020], or to both forearms [Liaci et al., 2016]. The position of each consecutive stimulus pair alternated between the opposing diagonal corners of an imaginary rectangle which leads to switches between the perception of motion travelling either horizontally or vertically. The proximity of stimulus pairs is the strongest contributor to the direction in which apparent motion is perceived, with motion more likely to be experienced between closer stimulus pairs than more distant ones [Gengerelli, 1948].

There are numerous stimulus examples for perceptual ambiguity which show similar properties across sensory modalities and across different paradigms within the same sensory modality. Levelt’s propositions have been broadly used to describe perceptual rivalry in the visual [Moreno-Bote et al., 2010, Brascamp et al., 2015] and auditory [Rankin et al., 2015] domains, for example the generalization of Levelt’s proposition II states that increasing the difference between percept strengths increases the mean perceptual dominance of the stronger percepts [Levelt, 1965]. Despite mean dominance times varying widely in multistable experiment, across different observers and stimulus contrasts [Zhou et al., 2004, Brascamp et al., 2005], the distribution of perceptual phases maintains a constant shape (gamma-like distribution) [Blake et al., 1980, Cao et al., 2016, Denham et al., 2018]. Previous studies in vision suggested that the durations of successive perceptual phases are statistically independent (insignificant correlation) [Logothetis et al., 1996b, Pressnitzer and Hupé, 2006], however, recent studies of binocular rivalry [van Ee, 2009, Cao et al., 2020] revealed positive correlations for perceptual phases that were one phase apart (between different percepts). Auditory streaming experiments also show positive correlations for durations separated by one phase and negative correlations for durations that are two phases apart (between same percepts) [Barniv and Nelken, 2015].

Levelt’s propositions and other common characteristics in multistable perception have yet to be explored with tactile stimuli. Here we present a new ambiguous tactile stimulus paradigm, similar to beta motion from vision, presented at only two locations on the skin but still capable of producing tactile rivalry during minutes-long trials. Whilst beta motion in vision results from a fixed intensity (contrast) moving dot, the stimulus presented here involves alternating changes in intensity (vibration amplitude) at the locations on each index finger. Two different perceptual interpretations can arise that compete for dominance over time. Because of the simplicity of this tactile stimulus, it is well suited to investigate the mechanisms underlying tactile rivalry. We varied intensity difference (Δ*I*) asymmetrically during different trials as a control parameter to assess its effect on perceptual durations. We aim to characterise the stochasticity of perceptual alternations by looking at properties of the distribution of switching times. Our results show that general characteristics of bistable perception such as Levelt’s proposition II (LVII) and stochastic characteristics like a scaling property hold for the tactile modality. This provides a new avenue to expand studies of the general mechanisms underlying neural competition across multiple sensory modalities.

## Methods

### Participants

Fifteen volunteers (8 male, mean age 29.07 ± 7.43 SD) were recruited from the University of Exeter. Each gave written informed consent and received minor monetary compensation for participating in a one-hour session. Participants were naive to the purpose of the study and did not self-declare any neurological or sensory disorders. Procedures were in compliance with guidelines for research with human subjects and approved by University of Exeter Research Ethics Committee.

### Experiment design and procedure

Participants sat in an acoustically isolated booth and attended to vibrotactile stimulators attached to their right and left index fingers. We used miniature vibrotactile electromagnetic solenoid-type stimulators (18 mm diameter, Dancer Design tactors, www.dancerdesign.co.uk) driven by a tactile amplifier (Dancer Design Tactamp) to deliver stimuli. Vibrotactile stimuli consisted of sequences of 400 ms high (H) and low (L) intensity 200 Hz vibratory pulses, each followed by a 400 ms silent interval (H-L-HL for the right hand and L-H-L-H for the left hand, “-” indicates the silent gap) (Figure 1(A)). The intensity of the L stimulus was Δ*I* below the intensity of the H stimulus on a logarithmic scale (*dB*). We chose the full-amplitude based on a value where differences in amplitude were noticeable in unambiguous perception. The voltages applied to the tactors for full-amplitude 200 Hz sinosoidal vibration was 3.38 v. To mask any unwanted low-intensity sound emitted by the vibrotactile stimuli, participants listened to pink noise played through noise-isolating headphones at a comfortable listening level. During a trial participants’ perception of the stimulus changed over time and they reported their current perceptual interpretation with key presses (sampled at 100 Hz) on a keyboard (Figure 1(B)). Participants perceived the stimulus as either one simultaneous pattern of vibration on both hands (SIM), or patterns of vibration that jumped from one hand to the other hand, giving a sensation of apparent movement (AM) (Figure 1(C)). Subjects were instructed to report their percepts passively and not try to hold one perceptual interpretation over another.

**Figure 1.**
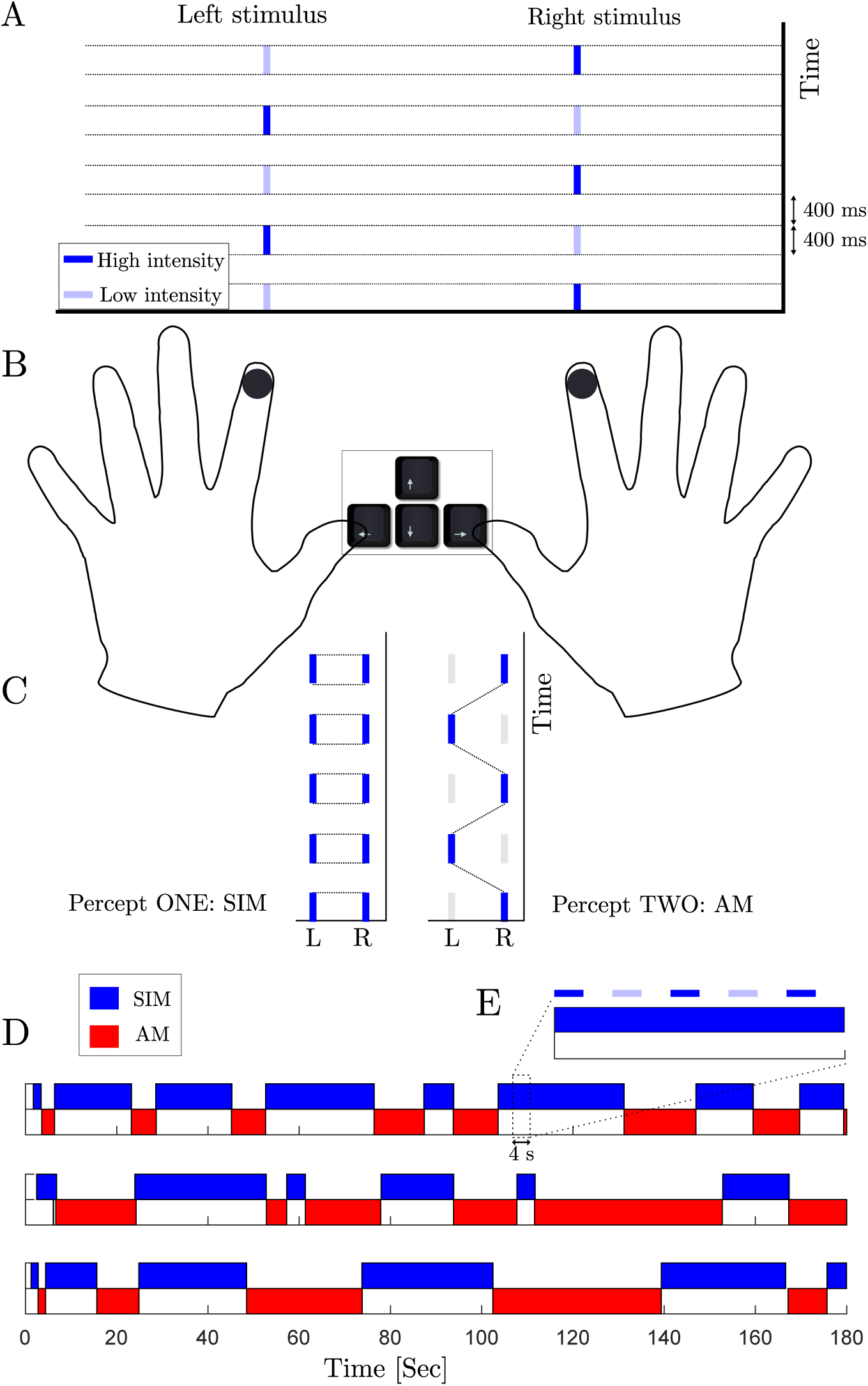
**(A) Vibrotactile stimuli.** Sequences of high (dark blue) and low intensity (light blue) of 200 KHz vibrations are delivered to the right and the left index finger. **(B) Experimental setup.** Vibrotactile stimuli are delivered to subject’s index fingers. During a trial subject’s perception of the stimuli changes and they report it by holding appropriate keys on the keyboard using their thumbs. **(C) percept types.** When the patterns are played with equal intensity, they can be perceived as one simultaneous vibration (SIM). With a fixed intensity difference (ΔI > 0 dB) between the high and low intensity tactile pulses, perception switches back and forth between two percepts: SIM (perceived as a fixed intensity on each hand, even though the intensity is changing) and AM (perceived as pulses of vibrations jumping from one hand to the other hand). We associate the left arrow key with SIM and the right arrow key with AM. **(D) Perceptual phases.** Perceptual interpretations of the stimuli for three different subjects during 3-minute trials at ΔI = 2 dB. **(E) Relative scale of the stimuli.** 4-second zoomed panel which shows the relation between the stimulus and the perceptual phases.

During experimental sessions, three-minute trials were repeated three times in blocks of 5 trials for a range of intensity differences (Δ*I* = 0.5, 1, 2, 4, 6 *dB*) in random order. For a given participant, this resulted in a total of 15 trials (45 minutes total trial time). The interval between trials was a minimum of 20 s. We used a 5 × 5 latin square design with nine randomized and unique grids so that the order of conditions for each participant in a block of trials was counterbalanced within/across participants and repetitions.

### Data analysis

In the first stage of our analysis, every trial with average percept durations larger than 150 s (above dashed line in Figure 2(A)) or smaller than 4 s (equivalent to H-L-H-L-H-) were excluded from data set before further analysis (91 trials were excluded from the total of 225 trials). For each of 15 subjects we take the average percept duration across all repetitions. So, each subject contributes a mean SIM and a mean AM duration averaged across all durations for the three repetitions at each condition. Other measures such as proportion of time with SIM or AM percept, and the frequency of switches are computed in a similar fashion. To see the effect of discarding trials with mean SIM and mean AM durations below 4 s and beyond 150 s compare Figure 2 with Figure S1 (no qualitative change).

**Figure 2.**
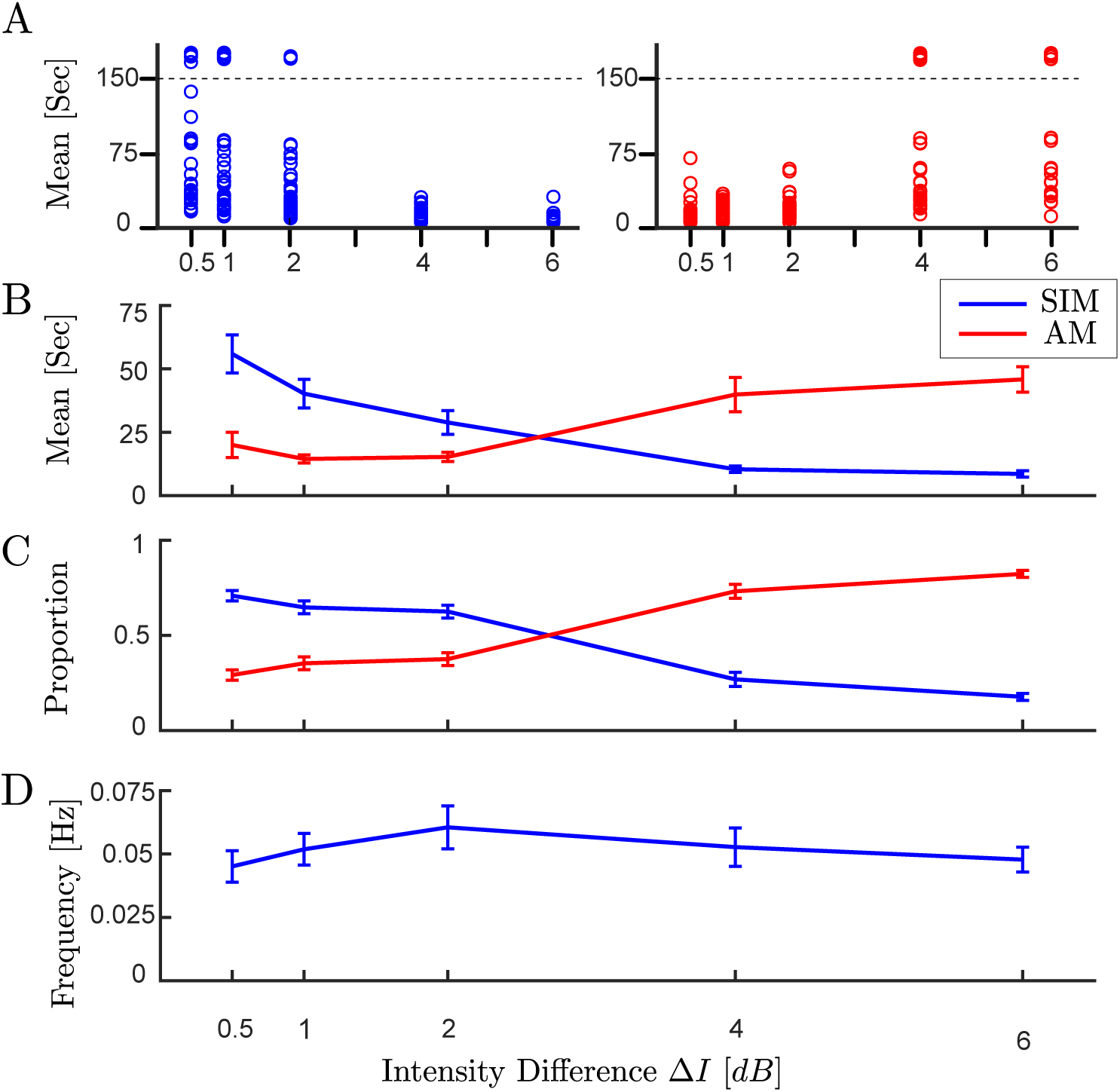
**(A)** The distribution of dominance durations at each trial with different experimental conditions (45 observations per experimental condition). All the samples above the dashed line (larger than 150 s), were excluded from the data set before further analysis (for similar analysis without excluding data see Figure S1 in supplementary materials). It is assumed that durations longer than 150 s are from trials with where perception did not alternate. **(B)** Mean dominance duration, **(C)** proportion of dominance for each percept type, **(D)** alternation rate, as a function of intensity difference (ΔI).

In order to establish whether the parameter Δ*I* had a significant effect on the measures described above, we used repeated measure ANOVA with Bonferroni corrections. As we had discarded some of the trials (due to unacceptably large or small means), we resampled data from mean percept durations at each experimental condition to substitute discarded trials and balance the number of observations across experimental conditions. ANOVA tables and posthoc analyses are reported in full in Table S1-S10 (supplementary material). A significance level of 0.05 is used throughout this paper. In ANOVA tables the Greenhouse-Geisser (G-G) corrected p-values are reported if a Mauchly sphericity (MS) test reached significance. In post-hoc analyses quoted p-values are Bonferroni corrected to account for multiple comparisons. Standard measures of effect size (generalized eta-squared) are quoted for statistically significant results. All statistical analyses were carried out in the statistical package R.

The distributions of normalized percept durations of both types shown in Figure 3 were compared with gamma and log-normal distributions using a one-way Kolmogorov-Smirnov (KS) test. The null hypothesis is that the test data are drawn from the standard distribution and a significant result (P < 0.05) indicates that the test data are not drawn from the comparison distribution.

**Figure 3.**
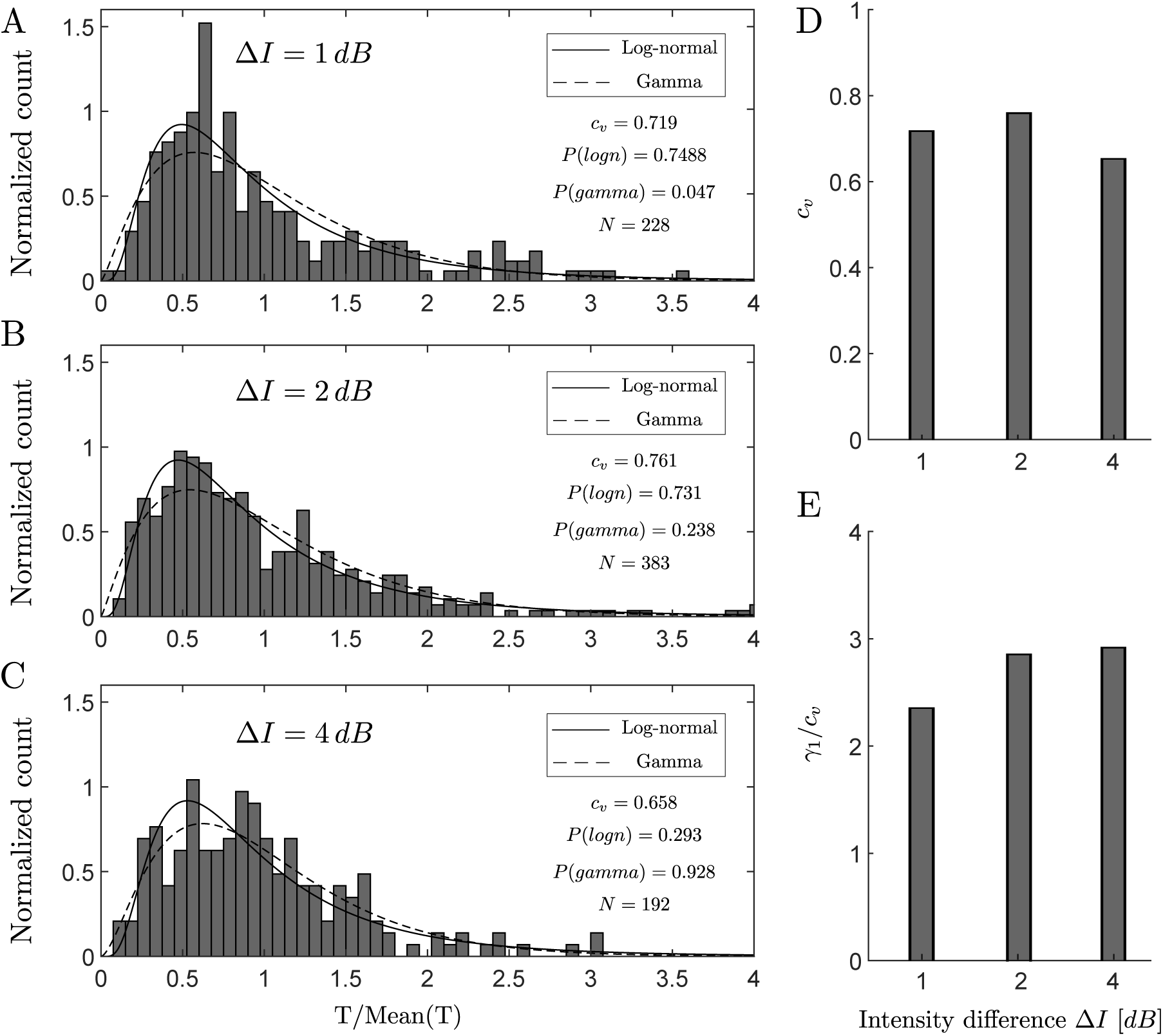
Histograms of normalized perceptual phases for experimental conditions close to equidominance, **(A)** ΔI = 1 dB, **(B)** ΔI = 2 dB and **(C)** ΔI = 4 dB combined across participants and percept type after normalization by the mean. Solid and dashed curves show the estimated log-normal and gamma distribution respectively. **(D)** Coefficient of variation (c_v_) and **(E)** skewness divided by coefficient of variation (γ_1_/c_v_) for experimental conditions ΔI = 1, 2, 4 dB.

In order to check for a scaling property, we first need to compute central moments. The distribution shape of samples t may be quantified in terms of the mean *µ*_1_ ≡ 𝔼 [*t*] and the central moments *µ*_2_ ≡ 𝔼 [(*t* − *µ*_1_)]^2^, *µ*_3_ ≡ 𝔼 [(*t* − *µ*_1_)]^3^, etc., or, equivalently, in terms of normalized moments, such as the coefficient of variation *c*_*v*_ = *µ*_2_^1/2^/*µ*_1_and the skewness *γ*_1_ = *µ*_3_/*µ*_2_^3/2^. A scaling property obtains if central moments are proportional to corresponding powers of the mean or, equivalently, if normalized moments are constant as follows:

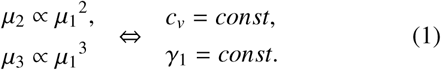

The normalization of percept durations was used in order to combine data across participants and different experimental conditions [Rankin et al., 2015], whilst accounting for some of the subject variability in experiments (supplementary material Figure S2). For correlation analysis, durations were normalized to the average value of each percept type separately within each trial and subject, in order to avoid spurious correlation due to inter-subject and intertrial differences in switching behaviour. Here we used the Pearson correlation that measures the strength and direction of the linear relationship between two variables. Pearson’s linear correlation coefficient *Corr* is defined as:

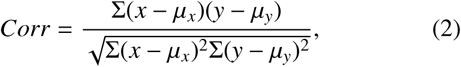

where *µ*_*x*_ and *µ*_*x*_ are the means of variables *x* and *y*.

## Results

The experiment with antiphase sequences of high and low intensity tactile stimuli on the right and the left index finger showed irregular perceptual switches between two interpretations for a range of the parameter Δ*I*. The average percept duration across all participants and trials for both percept types was 17.9 *s* ± 22.7 *s*, which spans on average over 35 periods of the stimuli (Figure 1D-E). The average percep duration across all participants for both percept types at Δ*I* = 2 *dB* was 15.8 *s* ± 19.4 *s*. The fraction of time spent in SIM was 0.52. For some participants at small or large Δ*I* values perception may not alternate, these trials were excluded from dataset.

### Levelt’s proposition II

To establish whether Levelt’s proposition II extends to the tactile domain, we chose intensity difference (Δ*I*) as a control parameter and examined the temporal dynamics of perceptual alternations. The distribution of mean perceptual dominance time in each trial across both percept types and Δ*I* conditions is plotted in Figure 2A. Mean dominance duration, proportion of time spent in each percept type and alternation rate are plotted against intensity difference (Δ*I*), respectively in Figure 2B-D. Increasing intensity difference (Δ*I*) (intensity of the H stimulus is fixed, intensity of the L stimulus is decreased), causes the mean dominance of SIM percept to decrease and AM percept to increase (Figure 2B). A similar pattern is shown for proportion of perceptual dominance (Figure 2C). The alternation rate reaches a maximum at equidominance Δ*I* = 2 *dB* (the point where each percept approximately dominates half the time), and decreases symmetrically below and above this point (Figure 2D). Of the conditions tested Δ*I* = 2 *dB* is closest to equidominance; from panels B and C in Figure 2 it appears Δ*I* = 3 *dB*, if tested, would be closer still to equidominance and have a higher alternation rate. Results from our experiments demonstrate that Levelt’s proposition II holds in tactile domain.

A two-way repeated measures ANOVA of mean duration of both percept types (SIM, AM) was performed with Percept type (SIM, AM) and intensity difference Δ*I* as within subjects factors. The analysis reported in Table 1 shows a significant interaction for Δ*I*:percept, *F*(4, 164) = 50.013, *P* < 10^−6^. As for the individual factor, percept does not reach significance *F*(1, 41) = 3.339, *P* = 0.075, however for the individual factor Δ*I* reaches significance, *F*(4, 164) = 3.929, *P* < 0.005. A one-way repeated measures ANOVA of mean duration of SIM percept was performed with Δ*I* as the within subjects factor. The analysis reported in Table 2 shows a significant effect of Δ*I* on mean dominance SIM *F*(4, 164) = 25.043, *P* < 10^−6^ (Similar results for AM percept in Table 4). Pairwise comparisons with Bonferroni-corrected significance levels revealed that each individual condition has significant differences for all non-adjacent conditions and for the pair Δ*I* = [2, 4] *dB* (Table 3, Similar results for AM percept in Table 5). Taken together, these result demonstrate a significant main effect of varying Δ*I* on mean dominance duration and proportion of dominance of each percept.

**Table 1.** Two-way repeated measure ANOVA of mean duration of **both percept types (AM, SIM)** with respect to intensity difference (ΔI) and percept type. Analysis shows a significant effect of ΔI and also ΔI:percept on the mean durations.

| Source | Type III SS | df | F | p | p < .05 | ges | p[G-G] |
| --- | --- | --- | --- | --- | --- | --- | --- |
| Percept | 1245.106 | 1 | 3.339 | 0.075 |  | 0.008 |  |
| Error(Percept) | 15288.68 | 41 |  |  |  |  |  |
| $\Delta I$ | 6553.935 | 4 | 3.929 | 0.004 | * | 0.038 | 0.009* |
| Error($\Delta I$) | 68380.52 | 164 | | | | | |
| $\Delta I$:Percept | 75498.04 | 4 | 50.013 | < $10^{-6}$ | * | 0.316 | < $10^{-6}$* |
| Error($\Delta I$:Percept) | 61892.01 | 164 | | | | | |

**Table 2.** One-way repeated measure ANOVA of mean duration of **SIM** perception with respect to intensity difference (ΔI). Analysis shows a significant effect of the intensity difference on the mean durations.

| Source | Type III SS | df | F | p | p< .05 | ges | p[G-G] |
| --- | --- | --- | --- | --- | --- | --- | --- |
| $\Delta I$ | 51582.3 | 4 | 25.043 | $< 10^{-6}$ | * | 0.029 | $< 10^{-6}$* |
| Error($\Delta I$) | 84450.65 | 164 | | | | | |

**Table 3.** Pairwise ttest, with Bonferoni corrected p-values, on the mean durations of **SIM** perception with respect to intensity difference (ΔI).

| | $\Delta I = 0.5$ | $\Delta I = 1$ | $\Delta I = 2$ | $\Delta I = 4$ |
| --- | --- | --- | --- | --- |
| $\Delta I = 1$ | 0.278 | - | - | - |
| $\Delta I = 2$ | 0.001 | 0.454 | - | - |
| $\Delta I = 4$ | $< 10^{-6}$ | $< 10^{-6}$ | 0.004 | - |
| $\Delta I = 6$ | $< 10^{-6}$ | $< 10^{-6}$ | 0.001 | 1.0 |

**Table 4.** One-way repeated measure ANOVA of mean duration of **AM** perception with respect to intensity difference (ΔI). Analysis shows a significant effect of the intensity difference on the mean durations.

| Source | Type III SS | df | F | p | p < .05 | ges | p[G-G] |
| --- | --- | --- | --- | --- | --- | --- | --- |
| $\Delta I$ | 30469.67 | 4 | 27.263 | $< 10^{-6}$ | * | 0.343 | $< 10^{-6}$* |
| Error($\Delta I$) | 45821.87 | 164 | | | | | |

**Table 5.** Pairwise ttest, with Bonferoni corrected p-values, on the mean durations of AM perception with respect to intensity difference (ΔI).

| | $\Delta I = 0.5$ | $\Delta I = 1$ | $\Delta I = 2$ | $\Delta I = 4$ |
| --- | --- | --- | --- | --- |
| $\Delta I = 1$ | 1.0 | - | - | - |
| $\Delta I = 2$ | 1.0 | 1.0 | - | - |
| $\Delta I = 4$ | $8.2 * 10^{-5}$ | $1.2 * 10^{-5}$ | $5.6 * 10^{-5}$ | - |
| $\Delta I = 6$ | $< 10^{-6}$ | $< 10^{-6}$ | $< 10^{-6}$ | 0.017 |

### Statistics of dominance durations and scaling property

Perceptual phases for bistable stimuli have been shown to be fit well by gamma or log-normal distributions. However, experiments with large numbers of participants found the log-normal distribution to be a better fit across visual and auditory bistable stimuli [Denham et al., 2018]. The distributions of normalized perceptual phases for experimental conditions close to equidominance are shown in Figure 3A-C. In the temporal analysis of perceptual durations for bistable stimuli, the perceptual phases are normalized by the mean for each percept type (see Figure S2 in supplementary material for normalization method). The coefficient of variation (*c*_*v*_), which is the ratio of the standard deviation to the mean, is used as a measure of variability in the perceptual phases. The result of one-way KS tests show that, except for the gamma standard distributions at Δ*I* = 1 *dB* (*P*(*gamma*) < 0.05), all the other histograms are compatible with the comparison distributions.

To assess how well tactile rivalry conforms to the scaling property reported in [Cao et al., 2014, Cao et al., 2016], we compared observations from three intermediate experimental conditions (Δ*I* = 1, 2, 4 *dB*). The extreme conditions were discarded from further analysis, as they had fewer perceptual phases leading to inaccurate computation of second and third moments. A scaling property obtains if normalized moments, such as the coefficient of variation *c*_*v*_ and ratio of skewness and coefficient of variation *γ*_1_/*c*_*v*_ are constant. Figure 3D-E illustrates the results in terms of the coefficient of variation and skewness across different experimental conditions. The coefficient of variation remained consistently near *c*_*v*_ = 0.6 (Figure 3D) and ratio of skewness and coefficient of variation *γ*_1_/*c*_*v*_ remained roughly constant. (Figure 3E). In other words, a scaling property was maintained over intermediate experimental conditions.

### Analysis of correlation

Figure 4A-D plots the normalized duration of each perceptual phase against the duration of the next, for the two possible transitions (lag1: SIM →AM and AM →SIM). Importantly, durations were normalized to the average value of each percept type separately within each trial and subject, in order to avoid spurious correlations due to inter-subject and inter-trial differences in switching behaviour (see Methods). For consecutive phase durations (lag 1), the correlation was small but significantly larger than zero (Figure 4A-B). For the phase durations that were one phase apart (lag 2: SIM →SIM and AM →AM), the correlation was significantly smaller than zero (Figure 4C-D).

**Figure 4.**
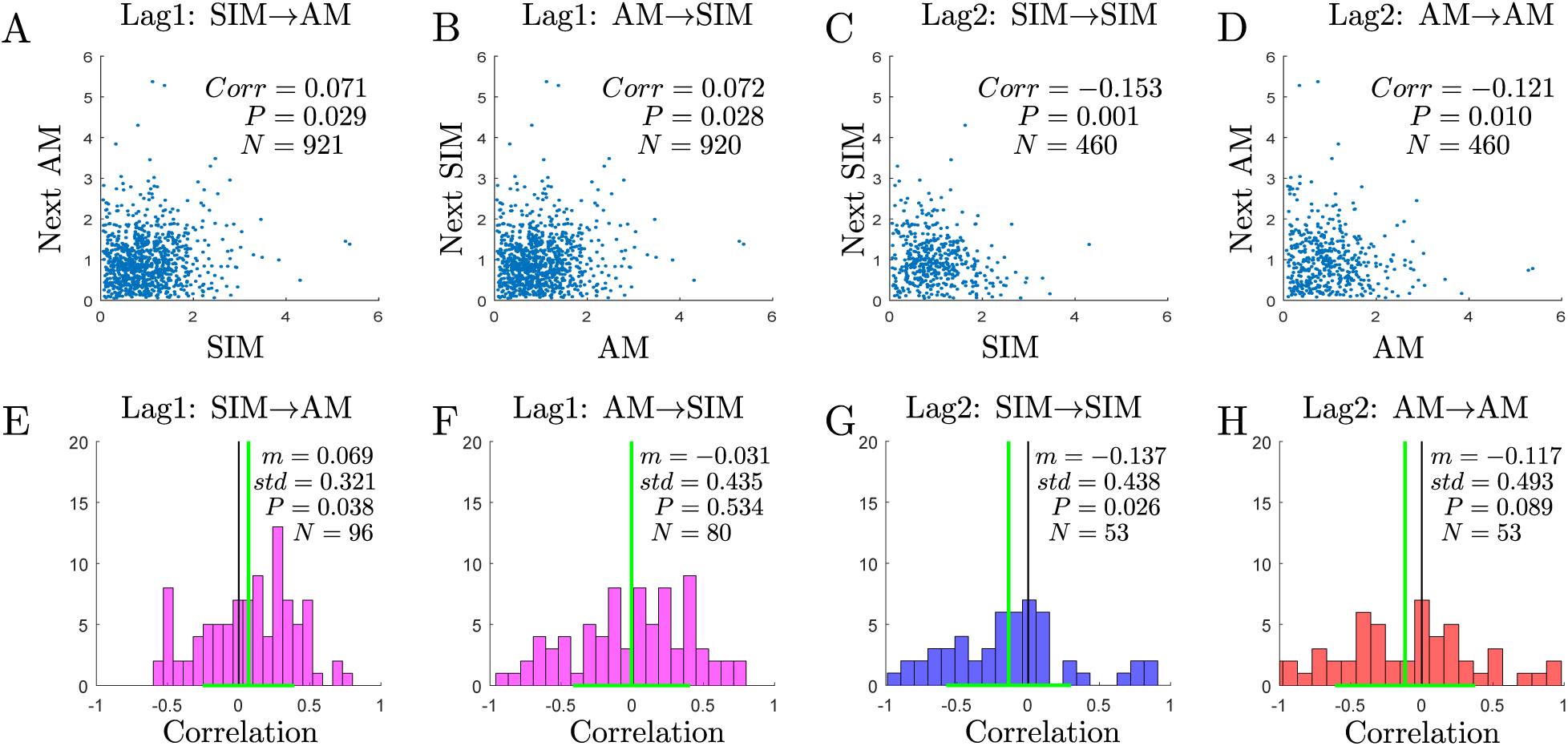
**(A-D)** Scatter plots of normalized durations. The correlation coefficient (corr) between perceptual phases for each scatter is indicated in each panel with the corresponding p-value and number of pairs (P and N respectively). **(E-H)** Histograms of correlation coefficients between perceptual phases in single trials. The mean (m) and standard deviation (std) are indicated in each panel, followed by the t-statistic of the fixed effect, its significance (P) and the number of trials for which it was possible to calculate the correlation (N). The black vertical lines mark zero correlation, and the green vertical lines mark the mean of each distribution. The transition types are marked above each histogram **(A**&**E)** AM following SIM. **(B**&**F)** SIM following AM. **(C**&**G)** SIM. **(D**&**H)** AM.

We also calculated correlation coefficients separately for each switch type in each individual trial. Figure 4E-H shows histograms of the single trial correlations between subsequent phases (lag 1) and between phase durations that are one phase apart (lag 2). Using a t-statistic, we found a significant deviation of the histogram towards positive correlation values for lag 1; SIM→AM transition (Figure 4E), and negative correlation values for lag2; SIM →SIM transition (Figure 4H). Single trial correlations for lag1; AM →SIM and lag 2; AM →AM were not significantly different from zero.

## Discussion

### Summary

Earlier studies with tactile stimuli have identified bistability in somatosensation; these have typically investigated the spatial proximity of stimulation sites as a control parameter. Future studies are needed to characterize the full range of spatial and temporal stimulation parameters capable of inducing bistable tactile apparent motion. Here we introduced a simple tactile stimulus which can evoke bistability through stimulating only two sites (in comparison with four sites in previous studies). We investigated the effect of varying intensity difference (Δ*I*) asymmetrically on perceptual durations. Our results show that Levelt’s proposition II (LVII) and other characteristics of sensory bistability that generalise in vision and auditory sensation extend to the tactile modality.

### Novelty of the introduced stimuli and experiment design

In this study we introduced a new ambiguous tactile stimulus, consisting of anti-phase sequences of high and low intensity vibration pulses on the right and left index fingertips. While in similar work on tactile bistability stimuli were presented on at least four sites on the skin [Carter et al., 2008, Liaci et al., 2016], our stimulus is simpler and was only presented at two sites. Our approach probes how motion perception is affected by feature (intensity) differences other than just location. This could be extended to look at other features e.g. vibration frequency, temporal cues, presentation rate, etc. Another important difference is in the way participants reported their perception and also in the analyses of perceptual responses. In one study, participants were asked to attend to the stimuli and report their perception at the end of short representations of tactile stimuli [Liaci et al., 2016]. In another study, participants reported their perception continuously during long representations of the stimuli, however; the analysis considered temporal evolution of the average responses pooled across all participants [Carter et al., 2008], which does not account for the dynamics of individual perceptual durations and inter-subject variability. In neither study is the intensity difference varied as a control parameter, but the results of our experiment show that it significantly affects the higher-level interpretation of the stimulus. Here, our analysis involves the temporal durations of perceptual phases and investigates the effect of varying the intensity difference (Δ*I*) asymmetrically on perceptual durations.

### Similar properties of perceptual competition across different modalities

Perception of the tactile stimuli showed the following similarities to other sensory modalities. First, visual, auditory and tactile stimuli can induce the perception of apparent motion which can be ambiguous in a certain range of stimulus parameters, for example here with a synchronous percept (SIM). Second, Levelt’s proposition generalizes to other visual modalities including ambiguous motion [Moreno-Bote et al., 2010] and to auditory bistability [Rankin et al., 2015]. Here our analysis showed Levelt’s proposition II extends to include tactile bistability. Third, in the experiments of visual and auditory bistability, the distribution of perceptual phases maintains a characteristic, gamma-like shape [Blake et al., 1980]. Denham et al. showed these distributions to be specifically log-normal in experiments with a large number of participants [Denham et al., 2018]. Even though mean perceptual phases vary widely between participants and stimulus properties, the variance and skewness of perceptual phases keep a characteristic proportion to the mean [Cao et al., 2014, Cao et al., 2016]. Our analysis shows that the scaling property holds in the tactile domain, similar to multistability in other modalities.

Despite these similarities, differences between the other sensory modalities and tactile perception were observed as well. Experiments with an auditory stimulus which utilised sequences of anti-phase high and low tones in each ear showed that the percept can be like a single tone oscillating from ear to ear, which is a form of apparent motion. However, there were other percepts like sensation of either the high tone in the left ear and the low tone in the right ear or vice versa, and the two percepts switching back and forth [Deutsch, 1974]. While we used a similar stimulus paradigm, the possible perceptual interpretations that we observed in the tactile domain were different. It remains to be determined whether perceptual interpretations equivalent to those reported in audition can be evoked by different ranges of spectrotemporal features in tactile stimuli. On the other hand, we found that perceptual phases in the tactile modality was generally more stable than in the other bistable perception like visual [Conrad et al., 2012] or auditory modality [Rankin et al., 2015] (mean dominance durations were longer in the tactile experiment). Previous studies in vision [van Ee, 2009, Cao et al., 2020] and auditory streaming [Barniv and Nelken, 2015] suggested that the durations of successive perceptual phases are positively correlated, however, for lag 2 they have been shown to be negatively correlated in auditory streaming (albeit their result did not prove to be statistically significant) [Barniv and Nelken, 2015] and to be positively correlated in binocular rivalry [Cao et al., 2020]. Our analysis with tactile stimuli shows a significant positive correlation for lag 1 (from one percept type to the other), and a negative correlation for lag 2 (between similar percept types).

### Locus of tactile rivalry and modelling

To the best of our knowledge, there are no models for tactile rivalry. Computational models of perceptual ambiguity have helped significantly with our understanding of other types of bistability. In order to develop a computational model of tactile rivalry, one must consider how inputs from the left and right hands project to primary somatosensory cortex (S1) and how features like amplitude, frequency and timing are encoded there. It is known that information from each half of the body-surface is projected to the opposite side of the brain [Maldjian et al., 1999]. Ipsilateral stimuli do not travel directly to the neurons in the primary somatosensory cortex (S1) [Eshel et al., 2010]. However, an extensive range of research from invasive studies in monkeys [Clarey et al., 1996], to behavioural or neuroimaging studies of humans [Hlushchuk and Hari, 2006] showed that somatosensory processing in the left side of the brain can be modulated by the right side and vice versa. For instance, during unilateral touch, contralateral activation can be observed as well as ipsilateral deactivation in S1 [Hlushchuk and Hari, 2006, Kastrup et al., 2008].

In contrast, intracranial recording [Noachtar et al., 1997], functional magnetic resonance imaging (fMRI) [Hämäläinen et al., 2002, Nihashi et al., 2005] and magnetoencephalography (MEG) [Korvenoja et al., 1995, Schnitzler et al., 1995] in humans, demonstrated that unilateral tactile stimulation activates bilateral S1 response. Moreover, single-cell recording in monkeys showed that some neurons in S1 display bilateral hand receptive fields [Iwamura et al., 2001]. These results suggest that interhemispheric connections may not be purely inhibitory for somatosensation and they are likely to have excitatory effects as well [Eshel et al., 2010]. We might use modelling to check whether purely inhibitory interactions can account for observed behavioural responses, or whether other types of interhemispheric interplay might explain the results instead.

The experiments presented here used tactile pulses at 200 Hz lasting 400 ms. The neural encoding of location and of spectro-temporal properties of tactile stimuli is relatively well documented. There exist quite a lot of discussions in the literature about the cortical representation of flutter frequencies (less than 80 Hz) [Talbot et al., 1968, Romo and Salinas, 2003], however, it is under debate whether the somatosensory cortices process high frequencies. There is some evidence to suggest that the temporal patterning of the neural activities in primary and secondary somatosensory cortex does not contain information about the frequency of the stimuli in the vibration range (greater than 80 Hz) [Ferrington and Rowe, 1980]. The perception of flutter and high frequency vibration may be processed through distinct processing streams [Tommerdahl et al., 2005]. Auditory cortex undoubtedly plays a main role in the spectro-temporal representation of acoustic stimuli. There are some hypotheses that the putative neural populations are independently stimulated by tactile and auditory sensory modalities [Yau et al., 2009]. The cutaneous vibration frequency can be distinguished at a significantly lower resolution (with Weber fractions around 0.2 for tactile versus 0.003 for auditory), and over a narrower range than auditory tone pitch (up to 1 kHz for tactile versus 20 kHz for auditory stimuli) [Saal et al., 2016]. The resolution of perception is likely hierarchical as supported by modelling work in visual [Wilson, 2003a] and auditory bistability [Rankin et al., 2015].

There is a need for a tactile rivalry model that accounts for both well-established results on the timing of dominance intervals and is also compatible with the physiological evidence and structure of somatosensation. General models of rivalry usually incorporate a slow process together with reciprocal inhibition to produce perceptual alternations. Perceptual bistability results from a competition between between units representing neural populations associated with different percepts (e.g. units driven by inputs from the left and right eyes in binocular rivalry) [Wilson, 2003b, Rankin et al., 2015, Cao et al., 2020]. A possible model of tactile rivalry could be based on competition between neural populations in primary somatosensory cortex associated with the right and the left hands. For the development of a model, an important aspect of tactile rivalry is that information is integrated across locations and over time to form the SIM and AM percepts. Dynamical analysis of models with different architecture and connectivity can help us to find a model that induces observed percept types with the specific characteristics and intrinsic dynamics. Our result show that Levelt’s proposition II (LVII) holds in the tactile modality. This provides a new avenue to test and expand the computational and neurobiological models for bistability currently dominated by vision science. The correlation structure and statistical distribution properties can be used as an important constraint on models of tactile rivalry, and could reveal mechanistic differences with other modalities.

## Conclusion

We presented a new, simple form of tactile rivalry and explored generalisations with respect to perceptual rivalry in other sensory modalities. First, the results of our study show that Levelt’s proposition II which describes the relation between stimulus features and mean perceptual dominance extends to tactile bistability. Second, the stochastic characteristics of irregular perceptual alternations were shown to follow similar distribution shapes across different experimental conditions and with different mean perceptual dominance (known as a scaling property). Third, in contrast with visual and auditory modalities, we found negative correlations for durations that were one phase apart (lag 2). The paradigm introduced here and the methods of analysis provide a basis to further explore how similar processes generate changes in perception across different senses. This opens up a new avenue for a range of experiment to explore the role of e.g.: adaptation and cross-inhibition in somatosensation, voluntary control (attention), perceptual memory and the integration of tactile cues with the other senses. This approach will allow for insights gained in previous work from a substantial literature on auditory and visual perception to deepen our understanding of tactile perception.

## Abbreviations

SIM: Simultaneo
AM: Apparent movement
H: High intensity
L: Low intensity
R: Right
L: Left

## Acknowledgements

We thank Alexander Billig, Raymond Van Ee, and John Rinzel for valuable feedback on earlier versions of this manuscript.

