## Supplementary materials for "Perceptual rivalry with vibrotactile stimuli"

### Supplementary material

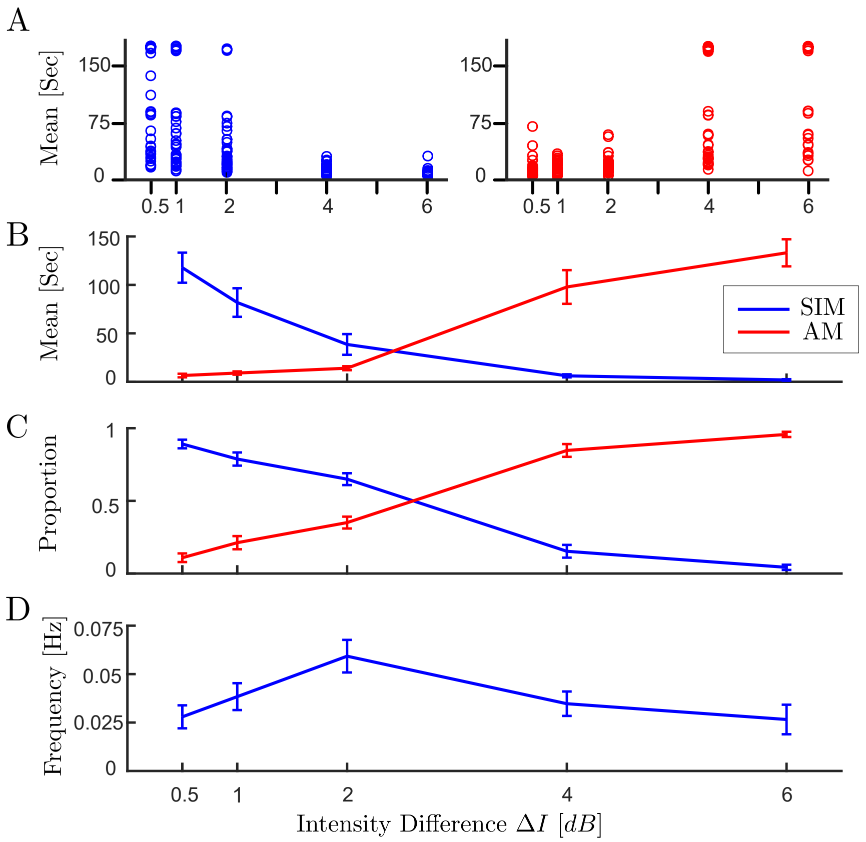


**Figure S1**

*Levelt's proposition II* **(A)** *The distribution of dominance durations at each trial with different experimental conditions (45 observation per experimental condition).* **(B)** *Mean dominance duration,* **(C)** *proportion of dominance for each percept type,* **(D)** *alternation rate, as a function of intensity difference ($\Delta I$).*

**Table S1**

*One-way repeated measure ANOVA of proportion of* ***both percept types (AM, SIM)*** *with respect to intensity difference [$\Delta I$] and percept type. Analysis shows a significant effect of $\Delta I$ and also $\Delta I$:percept on the proportion.*

| Source | Type III SS | df | F | p | p< .05 | ges | p[G-G] |
|---|---|---|---|---|---|---|---|
| Percept | 0.003 | 1 | 0.096 | 0.759 | | 0.001 | |
| Error(Percept) | 1.432 | 41 | | | | | |
| $\Delta I$ | 0.059 | 4 | 0.914 | 0.457 | | 0.006 | |
| Error($\Delta I$) | 2.648 | 164 | | | | | |
| $\Delta I$:Percept | 22.57 | 4 | 201.85 | $< 10^{-6}$ | * | 0.709 | $< 10^{-6}$* |
| Error($\Delta I$:Percept) | 4.584 | 164 | | | | | |

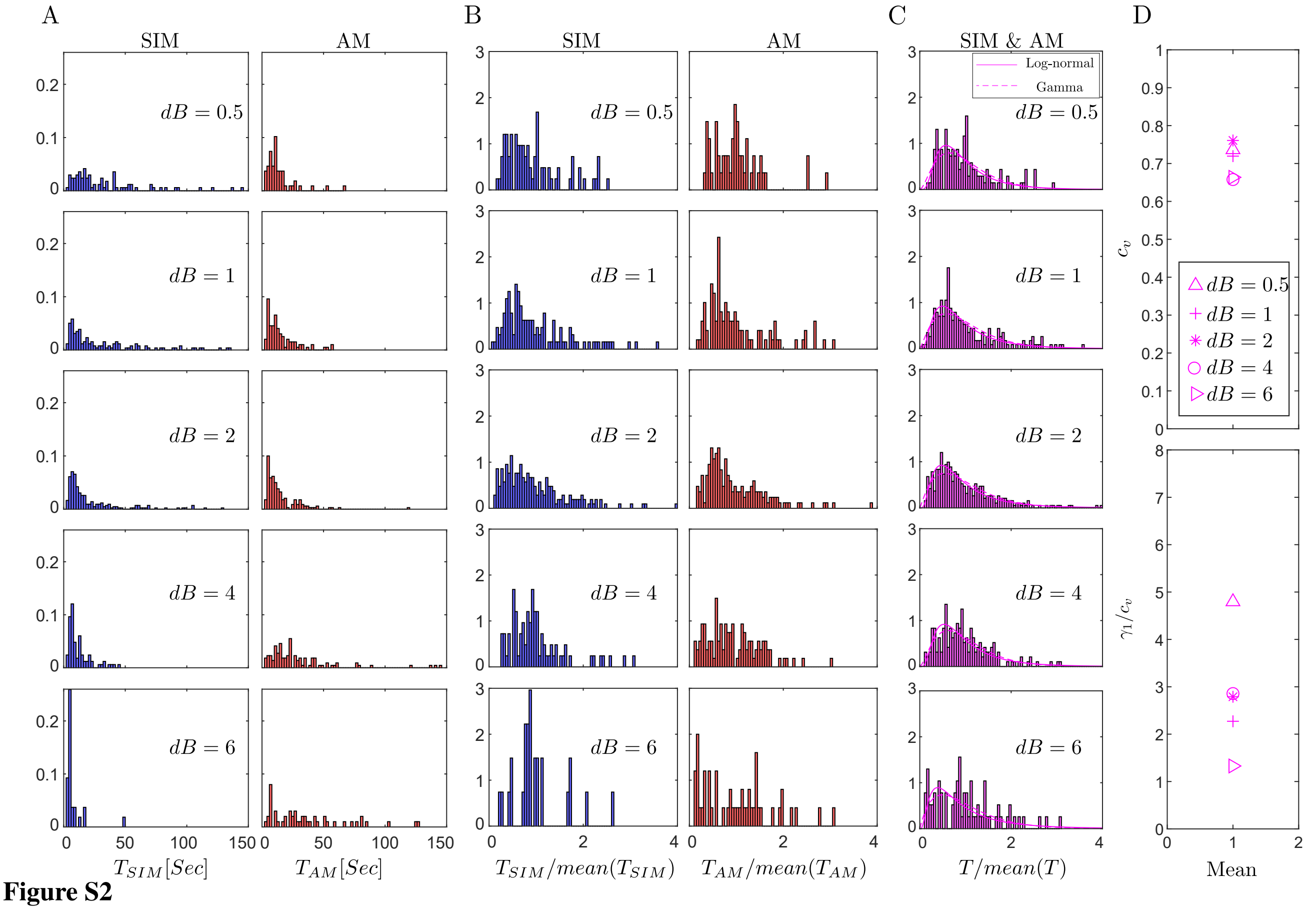


**Figure S2**

**(A)** *Histograms of perceptual phases for SIM (blue) and AM (red) percepts for each experimental condition.* **(B)** *Histograms of normalized perceptual phases. For normalization all samples of each histogram in panel (A) divided by its mean.* **(C)** *Normalized phase durations combined across the two percepts for each experimental condition. Solid and dashed curves show the estimated log-normal and gamma distribution respectively.* **(D)** *Coefficient of variation ($c_v$) and skewness divided by coefficient of variation ($\gamma_1/c_v$) for histograms in panel (C) with different experimental conditions ($\Delta I = 0.5, 1, 2, 4, 6\,dB$).*

**Table S2**

*One-way repeated measure ANOVA of proportion of* ***SIM*** *perception with respect to intensity difference ($\Delta I$). Analysis shows a significant effect of the intensity difference on the proportion.*

| Source | Type III SS | df | F | p | p< .05 | ges | p[G-G] |
|---|---|---|---|---|---|---|---|
| $\Delta I$ | 11.163 | 4 | 114.004 | $< 10^{-6}$ | * | 0.697 | $< 10^{-6}$* |
| Error($\Delta I$) | 4.014 | 164 | | | | | |

**Table S3**

*Pairwise ttest, with Bonferoni corrected p-values, on the proportion of **SIM** perception with respect to intensity difference ($\Delta I$).*

| | $\Delta I$ = 0.5 | $\Delta I$ = 1 | $\Delta I$ = 2 | $\Delta I$ = 4 |
|---|---|---|---|---|
| $\Delta I$ = 1 | 0.235 | - | | |
| $\Delta I$ = 2 | 0.001 | 0.454 | | |
| $\Delta I$ = 4 | $< 10^{-6}$ | $< 10^{-6}$ | $< 10^{-6}$ | |
| $\Delta I$ = 6 | $< 10^{-6}$ | $< 10^{-6}$ | $< 10^{-6}$ | 0.062 |

**Table S4**

*One-way repeated measure ANOVA of proportion of **AM** perception with respect to intensity difference ($\Delta I$). Analysis shows a significant effect of the intensity difference on the proportion.*

| Source | Type III SS | df | F | p | p< .05 | ges | p[G-G] |
|---|---|---|---|---|---|---|---|
| $\Delta I$ | 11.464 | 4 | 146.099 | $< 10^{-6}$ | * | 0.722 | |
| Error($\Delta I$) | 3.217 | 164 | | | | | |

**Table S5**

*Pairwise ttest, with Bonferoni corrected p-values, on the proportions of **AM** perception with respect to intensity difference ($\Delta I$).*

| | $\Delta I$ = 0.5 | $\Delta I$ = 1 | $\Delta I$ = 2 | $\Delta I$ = 4 |
|---|---|---|---|---|
| $\Delta I$ = 1 | 0.077 | - | | |
| $\Delta I$ = 2 | 0.005 | 1.00 | | |
| $\Delta I$ = 4 | $< 10^{-6}$ | $< 10^{-6}$ | $< 10^{-6}$ | |
| $\Delta I$ = 6 | $< 10^{-6}$ | $< 10^{-6}$ | $< 10^{-6}$ | $3 * 10^-5$ |

**Table S6**

*One-way repeated measure ANOVA of frequency of **both percept types (AM, SIM)** with respect to intensity difference ($\Delta I$) and percept type. Analysis shows a significant effect of $\Delta I$ and also $\Delta I$:percept on the frequency.*

| Source | Type III SS | df | F | p | p< .05 | ges | p[G-G] |
|---|---|---|---|---|---|---|---|
| Percept | 0.002 | 1 | 2.171 | 0.148 | | 0.004 | |
| Error(Percept) | 0.031 | 41 | | | | | |
| $\Delta I$ | 0.009 | 4 | 1.828 | 0.126 | | 0.020 | |
| Error($\Delta I$) | 0.205 | 164 | | | | | |
| $\Delta I$:Percept | 0.001 | 4 | 0.265 | 0.899 | | 0.002 | |
| Error($\Delta I$:Percept) | 0.124 | 164 | | | | | |

**Table S7**

*One-way repeated measure ANOVA of frequency of **SIM** perception with respect to intensity difference ($\Delta I$). Analysis shows a significant effect of the intensity difference on the frequency.*

| Source | Type III SS | df | F | p | p< .05 | ges | p[G-G] |
|---|---|---|---|---|---|---|---|
| $\Delta I$ | 0.006 | 4 | 1.846 | 0.122 | | 0.032 | |
| Error($\Delta I$) | 0.126 | 164 | | | | | |

**Table S8**

*Pairwise ttest, with Bonferoni corrected p-values, on the frequency of **SIM** perception with respect to intensity difference ($\Delta I$).*

| | $\Delta I$ = 0.5 | $\Delta I$ = 1 | $\Delta I$ = 2 | $\Delta I$ = 4 |
|---|---|---|---|---|
| $\Delta I$ = 1 | 1.00 | - | | |
| $\Delta I$ = 2 | 0.14 | 0.73 | | |
| $\Delta I$ = 4 | 1.00 | 1.00 | 1.00 | |
| $\Delta I$ = 6 | 1.00 | 1.00 | 0.71 | 1.00 |

**Table S9**

*One-way repeated measure ANOVA of frequency of AM perception with respect to intensity difference ($\Delta I$). Analysis shows a significant effect of the intensity difference on the frequency.*

| Source | Type III SS | df | F | p | p< .05 | ges | p[G-G] |
|---|---|---|---|---|---|---|---|
| $\Delta I$ | 0.004 | 4 | 0.859 | 0.489 | | 0.016 | |
| Error($\Delta I$) | 0.202 | 164 | | | | | |

**Table S10**

*Pairwise ttest, with Bonferoni corrected p-values, on the frequency of AM perception with respect to intensity difference ($\Delta I$).*

| | $\Delta I$ = 0.5 | $\Delta I$ = 1 | $\Delta I$ = 2 | $\Delta I$ = 4 |
|---|---|---|---|---|
| $\Delta I$ = 1 | 1 | - | | |
| $\Delta I$ = 2 | 0.84 | 1 | | |
| $\Delta I$ = 4 | 1 | 1 | 1 | |
| $\Delta I$ = 6 | 1 | 1 | 1 | 1 |
